# Probing the orientation specificity of excitatory and inhibitory circuitries in the primary motor cortex with multi-channel TMS

**DOI:** 10.1101/2021.08.20.457101

**Authors:** Victor Hugo Souza, Jaakko O. Nieminen, Sergei Tugin, Lari Koponen, Oswaldo Baffa, Risto J. Ilmoniemi

## Abstract

**Background:** The electric field orientation is a crucial parameter for optimizing the excitation of neuronal tissue in transcranial magnetic stimulation (TMS). Yet, the effects of stimulus orientation on the short-interval intracortical inhibition (SICI) and intracortical facilitation (ICF) paradigms are poorly known, mainly due to significant technical challenges in manipulating the TMS-induced stimulus orientation within milliseconds.

**Objective:** Our aim is to assess the effect of the TMS-induced stimulus orientation on the SICI and ICF paradigms and search for the optimal orientations to maximize the facilitation and suppression of the motor evoked potentials (MEP).

**Methods:** We applied paired-pulse multi-channel TMS in healthy subjects to generate SICI and ICF with conditioning and test pulses in the same, opposite, and perpendicular orientations to each other. The conditioning- and test-stimulus intensities were 80% and 110% of the resting motor threshold, respectively.

**Results:** Both SICI and ICF were significantly affected by the conditioning- and test-stimulus orientation. MEP suppression and facilitation were strongest with both pulses delivered in the same direction. SICI with a 2.5-ms and ICF with a 6.0-ms interstimulus interval (ISI) were more sensitive to changes in stimulus orientation compared with SICI at 0.5- and ICF at 8.0-ms ISIs, respectively.

**Conclusion:** Our findings provide evidence that SICI and ICF at specific ISIs are mediated by distinct mechanisms. Such mechanisms exhibit a preferential orientation depending on the anatomical and morphological arrangement of inhibitory and excitatory neuronal populations. We also demonstrate that the SICI and ICF can be maximized by adjusting the TMS-induced electric field orientation.

## Introduction

The inhibitory and excitatory neuronal circuits in the human neocortex are responsible for modulating the resulting cortical excitation, being critical for brain function. Paired-pulse transcranial magnetic stimulation (TMS) in the primary motor cortex can selectively assess the inhibitory and excitatory circuits by the short-interval intracortical inhibition (SICI) and intracortical facilitation (ICF) paradigms, respectively. In paired-pulse TMS, a conditioning stimulus is followed by a test stimulus at a millisecond-range interstimulus interval (ISI) [1]. Previous studies have demonstrated that a fine adjustment of the stimulation parameters, such as ISI, pulse shape, current direction and intensity, in both stimuli ultimately define the magnitude of the motor evoked potential (MEP) suppression or facilitation [2–6]. However, due to significant technical challenges in changing the TMS pulse orientation within milliseconds, little is known about how the stimulus orientation affect the excitatory and inhibitory in the living, intact human brain.

SICI and ICF are presumed to have separate neurophysiological origins for their distinct dependency on the stimulus parameters (for a review see Di Lazzaro and Rothwell (2014)). SICI has been attributed to emerge from two distinct mechanisms depending on whether the ISI is shorter or longer than 1 ms. First, an ISI shorter than 1 ms might inhibit the generation of descending volleys mainly by neuronal refractoriness due to the depolarization caused by a subthreshold conditioning stimulus [8,9]. Second, inhibition at ISIs between 1 and 5 ms is dominated by transsynaptic release of gamma-aminobutyric acid A (GABA_A_) [10–13]. In turn, with an ISI between 6 and 30 ms, motor responses are strengthened as a result of the ICF effect which seems to be mediated by N-Methyl-D-aspartic acid (NMDA) [14–16].

The underlying mechanisms and the neuronal populations associated with SICI and ICF phenomena play a critical role on the sensitivity of these phenomena to the orientation of the TMS-induced electric field. MEP facilitation has been shown to vanish when the conditioning stimulus is rotated from being across to being along the central sulcus in ICF protocols (ISIs of 6–20 ms), whereas suppression of motor response is maintained in SICI protocols (ISIs of 1–4 ms) [2]. Moreover, a TMS-induced current flowing in the anterior-posterior direction have been shown to induce smaller MEPs due to the strongest SICI compared with the posterior-anterior direction [5,17].

The neuronal morphology coupled with the orientation of the TMS-induced electric field in the cortical tissue is a determinant factor for the neural excitation [18,19]. Therefore, the dependency of SICI and ICF effects on the stimulus orientation is an indirect evidence of the structural-functional relationship of neuronal circuits in the primary motor cortex. For instance, GABA_A_ inhibitory interneurons mostly exhibit a stellate axonal arborization and excitatory interneurons usually have long-projected axonal branches across the cortical layers [20]. However, to our knowledge, there is no comprehensive assessment of how changes of the conditioning- and test-pulse orientations affect the magnitude of distinct SICI and ICF mechanisms.

The aim of this study was to describe the orientation sensitivity profile of distinct mechanisms engaged in SICI (0.5- and 2.5-ms ISI) and ICF (6.0- and 8.0-ms ISI). Furthermore, we estimated the optimal conditioning- and test-pulse orientation to maximize the SICI and ICF effects and selectively probe the inhibitory and excitatory neural circuits in the primary motor cortex. To deliver consecutive pulses in different orientations within the desired ISI, we employed multi-channel TMS (mTMS) technology [21], enabling an unprecedented approach to study the effect of stimulus orientation on the intracortical mechanisms at the time scale of neuronal excitation.

## Material and Methods

### Participants

Twelve healthy subjects (age: 29 years (mean), 27–41 years (range); 4 women) participated in the study. All participants were right-handed and gave written informed consent before their participation. The study was performed in accordance with the Declaration of Helsinki and approved by an Ethics Committee of the Hospital District of Helsinki and Uusimaa.

### Experimental procedure

Data were recorded in the same experimental sessions with another study [21], with shared subject preparation, motor mapping, and motor threshold determination. Briefly, surface electromyography (EMG) electrodes were placed over the right abductor pollicis brevis (APB), abductor digiti minimi, and first dorsal interosseous muscles in a belly–tendon montage. EMG was recorded with a Nexstim eXimia EMG device (500-Hz low-pass filtering, 3000-Hz sampling frequency; Nexstim Plc). The mTMS transducer was placed relative to the subject’s cortical anatomy using a neuronavigation system (NBS 3.2, Nexstim Plc) and an individual anatomical magnetic resonance image with voxel dimensions less than or equal to 1 mm. The hotspot and optimal orientation to elicit maximal MEP amplitudes was obtained with the electric field peak induced by the bottom coil being approximately perpendicular to the central sulcus and with the first phase of the induced peak electric field being anteromedial (AM). The resting motor threshold was defined as the lowest stimulus intensity capable of evoking at least 10 out of 20 MEPs with minimum peak-to-peak amplitude of 50 µV [22].

We employed a paired-pulse mTMS paradigm to evaluate the effect of stimulus orientation on inhibitory and excitatory mechanisms. The conditioning- and test-pulse intensities were 80 and 110% of the resting motor threshold in the AM orientation, respectively. We tested four ISIs (0.5, 2.5, 6.0, and 8.0 ms), four conditioning-pulse orientations (AM, posteromedial (PM), posterolateral (PL), and anterolateral (AL), and two test-pulse orientations (AM and PL). The orientation terminology refers to the direction of the peak induced electric field in the cortex. Twenty paired-pulses were administered for each combination of conditioning-pulse orientation, test-pulse orientation, and ISI. An additional 20 test pulses (single pulses) were applied in the AM and PL orientations, respectively, at an intensity of 110% of the resting motor threshold in the AM orientation to record reference MEPs.

The mTMS pulses had a trapezoidal monophasic waveform and were delivered by custom-made electronics [23,24]. The single-pulse stimulation intensity when determining the hotspot and the resting motor threshold was adjusted by varying the capacitor voltage with the current waveform phases lasting for 60.0 (rise), 30.0 (hold), and 43.2 µs (decay). For the paired-pulse stimulation, the desired intensities were obtained by manipulating the duration of the rise, hold, and decay current waveform phases, as described in detail by Nieminen et al. (2019). The waveform durations were defined based on a model of neuronal depolarization during the mTMS pulses, accounting for the capacitor voltage drop due to the conditioning pulse and a reference neuronal membrane time constant of 200 µs. This approach allowed us to apply a pair of pulses with a millisecond-scale interval while producing the desired stimulation effects in the cortical tissue. The conditioning-pulse phase durations were 43.8, 30.0, and 32.5 µs, and the test-pulse phase durations were 75.1, 30.0, and 52.7 µs. The paired-pulses order was pseudo-randomized and divided into 10 sequences containing 68 pulse pairs each, followed by short breaks of 2 to 5 min (including the time for cooling the transducer). The inter-train interval was pseudo-randomized from 4 to 6 s. The transducer temperature was monitored with a thermal infrared camera (FLIR i3, FLIR Systems, USA). If the surface temperature reached 41 °C, the transducer was cooled to about 30 °C.

### Data analysis

#### Preprocessing

MEPs were extracted from the EMG recordings. Trials showing muscle pre-activation or movement artifacts greater than ±15 µV within 200 ms before the TMS pulse were removed from the analysis. For each trial, we computed the MEP peak-to-peak amplitude at 15–60 ms after the TMS pulse and manually annotated the MEP latency. The latencies from trials with a peak-to-peak amplitude below 50 µV were rejected from the analysis due to low signal-to-noise ratio and increased uncertainty in accurately defining the onset time. In total, 1.2% of the trials were rejected and 28.3% of the MEP latencies were not annotated. The relatively high amount of rejected latencies was due to the small MEPs that occurred in the SICI protocols, as expected (see Results below). Data were pre-processed using custom-made scripts written in MATLAB R2021a (The MathWorks, Inc., USA).

#### Statistical analysis

The MEP amplitude and latency distributions were inspected visually for each subject and experimental condition to ensure a symmetric data distribution and similar variance across all conditions. The data of one subject were excluded from further analysis due to substantially small MEPs for all ISIs, consistently deviating from the effects observed in all other participants. The MEP amplitudes were log-transformed to correct for heteroskedasticity, i.e., variance inequalities across conditions [25]. The effect of the conditioning-pulse orientation, test-pulse orientation, and ISI on MEP amplitude and latency were assessed with linear mixed-effects models [26]. The 20 test pulse reference measurements were appended to the dataset to allow identification of inhibition and facilitation. They were replicated for all ISIs to maintain the balanced design and allow multiple comparisons between all conditions and the reference samples. The model comprised each condition and all possible interactions as fixed effects and a random structure with subject identifiers and correlated random intercepts and slopes for conditioning- and test-pulse orientations. The random structure was selected based on sequential testing of hierarchical modeling with each model fit using likelihood-ratio tests [27,28]. The selected model was recomputed using restricted maximum likelihood estimation and *p*-values for fixed-effects derived with Satterthwaite approximations in a Type III Analysis of Variance table. Post-hoc multiple comparisons were computed with estimated marginal means and p-values corrected for false discovery rate [29]. The model diagnostic was performed with Q–Q plots of residuals to assess any critical deviation from the normal distribution, and standard versus fitted values plots to inspect for heteroscedasticity. Statistical analysis was performed using custom-made scripts written in R 3.6 (R Core Team, Vienna, Austria) with the *lme4* 1.1 and *afex* 0.25 packages for computing the linear mixed-effects models, and *emmeans* 1.4 for computing the estimated marginal means. The level of statistical significance was set to 0.05.

## Results

The effects of test-pulse orientation, conditioning-pulse orientation, and ISI on the MEP amplitude and latency are illustrated in Figure 1. The MEP amplitude inhibition and facilitation were significantly affected by changes in the test- and conditioning-pulse orientation and ISI, as suggested by the significant interaction term (TP × CP × ISI) in Table 1. The MEP latency also varied depending on the pulse orientations. All multiple comparison results are shown in Supplementary Tables 1–6.

**Figure 1:**
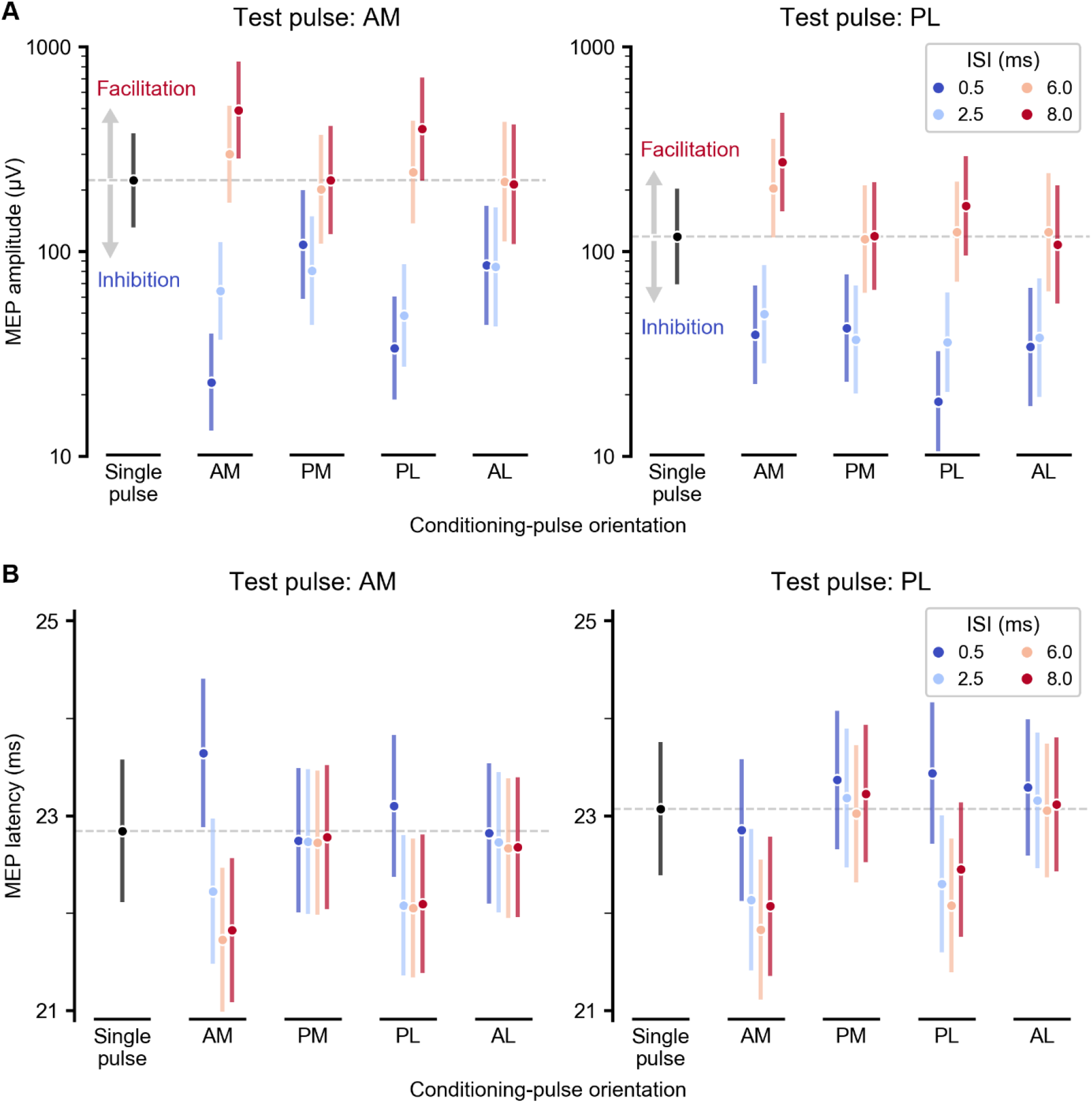
MEP amplitude (**A**) and latency (**B**) as a function of the conditioning-pulse orientation and interstimulus interval (ISI) for the test pulse in the anteromedial (AM, left) and posterolateral (PL, right) orientations. The circular symbols mark the MEP amplitude or latency estimated by the linear mixed-effects model, and the vertical bars indicate the corresponding 95% confidence interval of the mean. The horizontal dashed light gray line indicates the single-pulse MEP amplitude (**A**) or latency (**B**) in the model. The conditioning pulses were delivered in the AM, posteromedial (PM), PL, and anterolateral (AL) orientations.

**Table 1:** Type III Analysis of Variance Table with Satterthwaite’s method for the linear mixed-effects model of the MEP amplitude. Fixed factors were the test-pulse orientation (TP), conditioning-pulse orientation (CP), and interstimulus interval (ISI). Interaction between factors is represented as “×”.

| Effect | Degrees of freedom<br>(numerator, denominator) | F-value | p-value |
| --- | --- | --- | --- |
| TP | (1, 10.0) | 28.38 | < 0.001 |
| CP | (4, 12.2) | 7.96 | 0.002 |
| ISI | (3, 8578.2) | 769.20 | < 0.001 |
| TP × CP | (4, 8578.2) | 17.47 | < 0.001 |
| TP × ISI | (3, 8578.2) | 2.08 | 0.100 |
| CP × ISI | (12, 8578.2) | 80.77 | < 0.001 |
| TP × CP × ISI | (12, 8578.2) | 6.57 | < 0.001 |

**Table 2:** Type III Analysis of Variance Table with Satterthwaite’s method for the linear mixed-effects model of MEP latency. Fixed factors were the test-pulse orientation (TP), conditioning-pulse orientation (CP), and interstimulus interval (ISI). Interaction between factors is represented as “×”.

| Effect | Degrees of freedom<br>(numerator, denominator) | F-value | p-value |
| --- | --- | --- | --- |
| TP | (1, 10.9) | 18.56 | 0.001 |
| CP | (4, 12.8) | 16.81 | < 0.001 |
| ISI | (3, 5688.5) | 79.93 | < 0.001 |
| TP × CP | (4, 5544.0) | 14.83 | < 0.001 |
| TP × ISI | (3, 5605.5) | 2.15 | 0.092 |
| CP × ISI | (12, 5399.8) | 20.03 | < 0.001 |
| TP × CP × ISI | (12, 5689.8) | 3.00 | < 0.001 |

### Effect of stimulus orientation on SICI

We observed different levels of MEP suppression depending on the conditioning- and test-pulse orientation at 0.5- and 2.5-ms ISIs. With the test pulse in the AM orientation and at a 0.5-ms ISI, the AM conditioning pulse caused the strongest MEP inhibition (smallest amplitudes) compared to the other orientations. Conditioning pulses in the PM and AL orientations, i.e., perpendicular to the test pulse, generated considerably less MEP suppression (higher amplitudes) than conditioning pulses in the AM and PL orientations. For instance, an AM conditioning-pulse orientation elicited MEP amplitudes about 9.7 times smaller than the test pulse alone, compared with an amplitude reduction of 6.6, 2.6, and 2.0 times for conditioning pulses in PL, AL, and PM directions, respectively (for all comparisons *p* < 0.001). The effect of conditioning-pulse orientation was different for the 2.5-ms and 0.5-ms ISI. For the 2.5-ms ISI, a PL conditioning pulse evoked the smallest MEP amplitudes (conditioning and test pulses in opposite directions) compared with the AM and PM orientations; conditioning pulses in PL and AL orientations resulted in similar amplitudes. These changes in the level of MEP suppression were less prominent with the test pulse in the PL orientation. The pulse configuration showing strongest MEP suppression (AM–AM and PL–AM at 0.5-ms ISI) also generated MEP latencies longer than the single-pulse MEPs. Interestingly, with a pair of pulses in AM–AM orientation, the latencies at all other ISIs were 1–1.5 ms shorter than with the test pulse alone. Conditioning pulses in the PM or AL orientation yielded MEPs with similar latencies than single-pulse MEPs regardless of the ISI and the test-pulse orientation.

### Effect of stimulus orientation on ICF

MEP facilitation was observed only when the test and conditioning pulses were either in the same or opposite directions and vanished for the conditioning stimulus inducing an electric field perpendicular to the test pulse. With an 8.0-ms ISI, the conditioning and test pulses in AM and PL orientations facilitated the MEP amplitude, whereas with a 6.0-ms ISI only the AM-oriented conditioning pulse resulted in facilitation. The shortest MEP latencies were obtained at 6.0- and 8.0-ms ISIs with the conditioning and test pulses in the same or opposite directions. When the conditioning and test pulses were perpendicular to each other, there was no significant difference in latency compared to the latency of single-pulse MEP at any ISI.

## Discussion

Our results provide evidence that cortical inhibition and facilitation exhibit a more complex dependence on the conditioning-stimulus orientation than previously reported and that these phenomena seem to be mediated by distinct mechanisms within the motor cortex. We demonstrated that the level of MEP amplitude suppression following the SICI protocols depended on the conditioning-stimulus orientation. Moreover, MEP potentiation occurred only when the conditioning stimulus was delivered in the same or opposite direction as the test stimulus.

Our findings showed that a subthreshold conditioning stimulus followed by a suprathreshold test stimulus, either 0.5 or 2.5 ms later, caused a suppression in MEP amplitude in all the tested conditioning-pulse orientations. Strikingly, the amount of inhibition generated from a 0.5-ms ISI significantly depended on the conditioning-stimulus orientation. Inhibition was strongest when the conditioning and test stimulus were given in the same or opposite directions and slightly reduced when the conditioning pulse was delivered at PM and AL orientations. We devised two possible explanations for such observations. First, ISIs may originate from the combination of axonal refractoriness and activation of inhibitory neuronal populations in superficial layers. Recently, Kurz et al. (2019) studied the activation of different layers in the motor cortex by TMS and electrical stimulation. According to that study, the I1-wave seems to have two components delayed 0.6 ms from each other. The later components may be generated on superficial layers 2/3, while the earlier from direct activation of pyramidal neurons in layers 5 [30]. In this sense, a subthreshold conditioning TMS pulse may activate superficial layers that inhibit the subsequent activation of deeper layers by the suprathreshold test pulse. It has already been demonstrated that inhibitory mechanisms have a considerably lower threshold for activation than the motor threshold itself [1,31–33]. Furthermore, the 0.6 ms closely coincides with the 0.5-ms ISI that showed the strongest inhibition in our study. Thus, our study may provide evidence to support the findings of Kurz et al. (2019). Interestingly, GABA_A_-mediated inhibitory interneurons project mainly horizontally within layers 2/3 with a distributed arborization [34,35]. Like the excitatory mechanisms, a set of neuronal projections running from layers 2/3 to 5 would be optimally activated by a stimulus in the AM or PL orientations, and explain the lower inhibition observed when conditioning pulse was delivered in the AL or PM orientations with a 0.5-ms ISI. Second, the conditioning stimulus may induce refractoriness in specific segments of pyramidal neurons that are orientation sensitive. In this case, the subsequent test stimulus would be less effective in directly stimulating those sites. However, it seems unlikely that both mechanisms would co-exist. Importantly, the stimulation intensity was adjusted for all conditions relative to the AM-orientation resting motor threshold; if otherwise adjusted for each orientation would aid differently in distinguishing the underlying mechanisms. Either way, our findings demonstrate that the predominant inhibitory mechanism seems to be non-specific to the stimulus orientation.

MEP facilitation was mainly observed when the conditioning stimulus was in the AM orientation, for 6.0- and 8.0-ms ISIs and both AM and PL test-pulse orientations. Such specificity possibly indicates that ICF is mediated by connections across layers within a cortical column and thus exhibits a higher sensitivity to changes in the induced electric field orientation in the cortical tissue. Indeed, excitatory neurons exhibit a preferential direction projecting from layers 2/3 to pyramidal neurons in layer 5 [34,36], and thus might not be excited by the conditioning pulses in the AL and PM orientations. The subthreshold conditioning pulse possibly activated the low-threshold, fast inhibitory interneurons which were later (after about 5 ms) depressed [37], leaving the cortical circuitry in an excited state. The increase in cortical excitability for the conditioning AM and PL pulses might also explain the observed shorter MEP latencies for 6.0-ms and 8.0-ms compared to the single-pulse stimulation. The excitation of intralayer projections evidences a similar orientation-dependency than that of the single-pulse MEPs, in which a stimulus in the AM orientation induces significantly higher neuronal excitation than in the PL orientation, as reported above. Interestingly, the longer 8.0-ms ISI potentiated the MEPs even for conditioning stimulus in the PL orientation. The additional delay might contribute further to a reduction of existent GABA inhibition [37], and thus allow stronger neuronal activation even with a stimulus in the suboptimal PL orientation. The origin of mechanisms underlying MEP facilitation on ICF protocols is a composition of cortical [38] and subcortical mechanisms [39,40]. The absence of ICF effects in the amplitude of descending volleys and existing facilitation of spinal reflexes by the conditioning pulse may suggest that facilitation has a substantial component at a subcortical level [40]. In turn, the modulation of transcranial-evoked responses evidences the primary enrolment of cortical mechanisms [38]. Our results provide compelling data on the direct relation between the orientation sensitivity of intracortical mechanisms and the MEP facilitation, further describing the structural and physiological principles of the mechanisms mediating ICF at the cortical level.

The effect of pulse orientation on SICI and SICF may also be influenced by the reduced neuronal excitation generated by pulses oriented along the central sulcus (AL and PM orientations). In a similar study from our group, we observed that adjusting the intensity of the conditioning pulse relative to the orientation-specific MT led to a stronger MEP suppression with perpendicular rather than same-direction conditioning and test pulses [41]. This finding was most likely due to the 33% difference between the two conditioning stimulus intensities. In fact, changes in the stimulus orientation are essentially a variation in the effective stimulation intensity, and these two parameters are physically connected. Therefore, the interaction between stimulus orientation and intensity must be carefully considered to maximize the SICI and ICF effects.

## Acknowledgments

This research has received funding from the Academy of Finland (Decisions No. 255347, 265680, 294625, and 306845), the Finnish Cultural Foundation, Jane and Aatos Erkko Foundation, Erasmus Mundus SMART2 (No. 552042-EM-1-2014-1-FR-ERA MUNDUSEMA2), the Conselho Nacional de Desenvolvimento Científico e Tecnológico (CNPq; grant number 140787/2014-3), the European Research Council (ERC) under the European Union’s Horizon 2020 research and innovation programme (grant agreement No 810377), and the Research, Innovation and Dissemination Center for Neuromathematics (FAPESP; grant number 2013/07699-0).

## Supplementary Material

**Supplementary Table 1:**
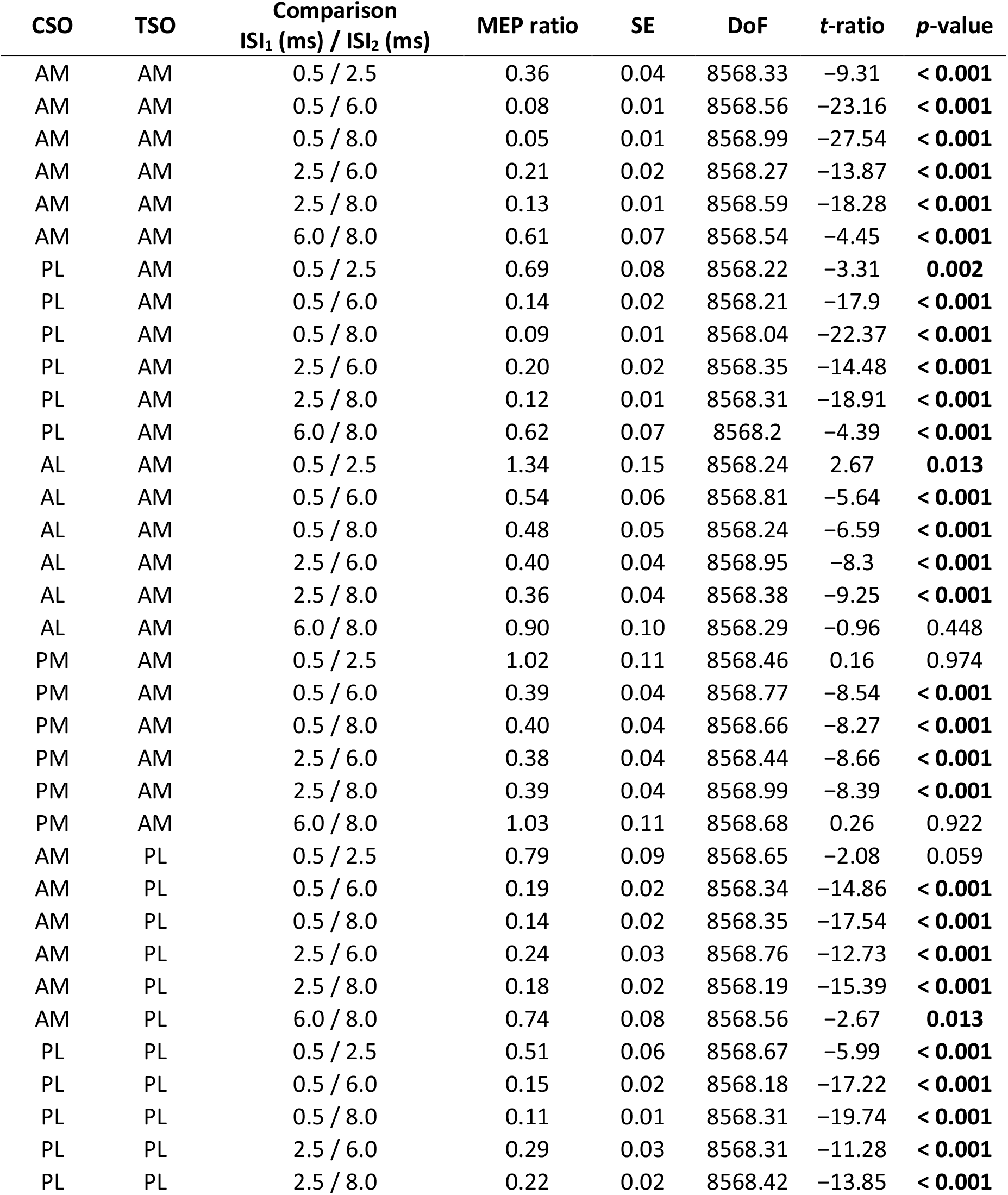
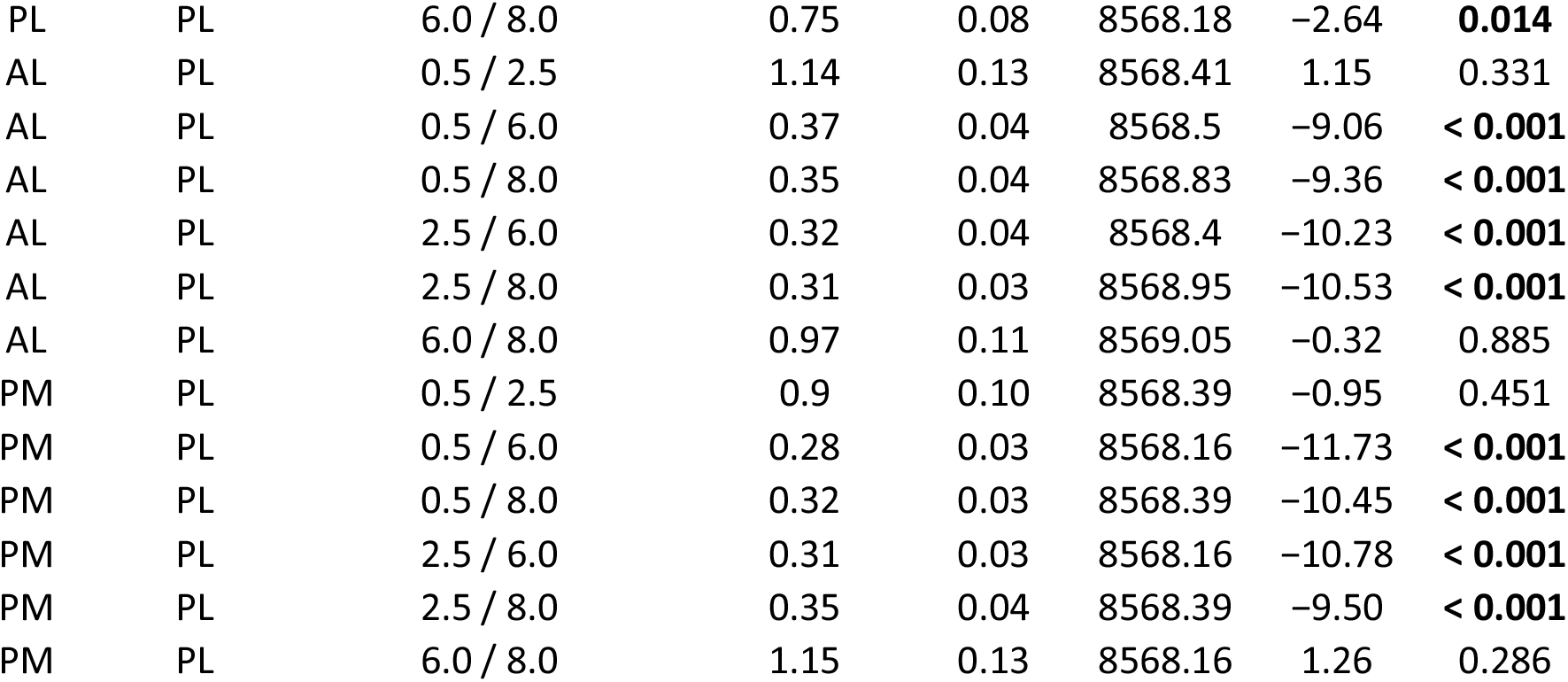
Multiple comparisons between interstimulus intervals (ISI). The results are presented as follows, for a fixed conditioning-stimulus orientation (CSO) and test-stimulus orientation (TSO), we compute the motor evoked potential (MEP) amplitude ratio (MEP ratio) between the tested ISI. Each comparison has a standard error (SE), degrees of freedom (DoF), *t*-ratio, and *p*-value. Tested stimulus orientations were anteromedial (AM), posterolateral (PL), anterolateral (AL), and posteromedial (PM). *p*-values in bold are smaller than the statistical significance level of 0.05.

**Supplementary Table 2:** Multiple comparisons between conditioning-stimulus orientations (CSO). The results are presented as follows, for a fixed test-stimulus orientation (TSO) and interstimulus interval (ISI), we compute the motor evoked potential (MEP) amplitude ratio (MEP ratio) between the tested CSO. Each comparison has a standard error (SE), degrees of freedom (DoF), *t*-ratio, and *p*-value. Tested stimulus orientations were anteromedial (AM), posterolateral (PL), anterolateral (AL), and posteromedial (PM). *p*-values in bold are smaller than the statistical significance level of 0.05.

| TSO | ISI (ms) | Comparison<br>CSO<sub>1</sub> / CSO<sub>2</sub> | MEP ratio | SE | DoF | *t*-ratio | *p*-value |
| --- | --- | --- | --- | --- | --- | --- | --- |
| AM | 0.5 | SP / AM | 9.69 | 1.52 | 30.85 | 14.48 | **< 0.001** |
| AM | 0.5 | SP / PL | 6.60 | 1.13 | 24.32 | 10.99 | **< 0.001** |
| AM | 0.5 | SP / AL | 2.06 | 0.34 | 26.28 | 4.34 | **< 0.001** |
| AM | 0.5 | SP / PM | 2.61 | 0.51 | 19.06 | 4.88 | **< 0.001** |
| AM | 0.5 | AM / PL | 0.68 | 0.09 | 70.56 | -2.95 | **0.007** |
| AM | 0.5 | AM / AL | 0.21 | 0.04 | 23.32 | -8.82 | **< 0.001** |
| AM | 0.5 | AM / PM | 0.27 | 0.05 | 21.08 | -7.11 | **< 0.001** |
| AM | 0.5 | PL / AL | 0.31 | 0.05 | 29.93 | -7.34 | **< 0.001** |
| AM | 0.5 | PL / PM | 0.39 | 0.06 | 26.96 | -5.65 | **< 0.001** |
| AM | 0.5 | AL / PM | 1.26 | 0.15 | 158.28 | 1.97 | 0.077 |
| AM | 2.5 | SP / AM | 3.47 | 0.54 | 30.92 | 7.93 | **< 0.001** |
| AM | 2.5 | SP / PL | 4.58 | 0.79 | 24.6 | 8.83 | **< 0.001** |
| AM | 2.5 | SP / AL | 2.77 | 0.46 | 26.38 | 6.11 | **< 0.001** |
| AM | 2.5 | SP / PM | 2.65 | 0.52 | 19.17 | 4.96 | **< 0.001** |
| AM | 2.5 | AM / PL | 1.32 | 0.17 | 72.22 | 2.11 | 0.059 |
| AM | 2.5 | AM / AL | 0.80 | 0.14 | 23.45 | -1.28 | 0.288 |
| AM | 2.5 | AM / PM | 0.76 | 0.14 | 21.26 | -1.46 | 0.223 |
| AM | 2.5 | PL / AL | 0.61 | 0.10 | 30.47 | -3.15 | **0.006** |
| AM | 2.5 | PL / PM | 0.58 | 0.10 | 27.52 | -3.30 | **0.005** |
| AM | 2.5 | AL / PM | 0.96 | 0.11 | 162.05 | -0.37 | 0.853 |
| AM | 6.0 | SP / AM | 0.75 | 0.12 | 31.14 | -1.87 | 0.105 |
| AM | 6.0 | SP / PL | 0.91 | 0.16 | 24.5 | -0.53 | 0.745 |
| AM | 6.0 | SP / AL | 1.11 | 0.18 | 26.17 | 0.61 | 0.685 |
| AM | 6.0 | SP / PM | 1.02 | 0.20 | 19.06 | 0.08 | 1.000 |
| AM | 6.0 | AM / PL | 1.22 | 0.16 | 72.45 | 1.55 | 0.177 |
| AM | 6.0 | AM / AL | 1.49 | 0.26 | 23.41 | 2.25 | 0.054 |
| AM | 6.0 | AM / PM | 1.36 | 0.25 | 21.22 | 1.67 | 0.156 |
| AM | 6.0 | PL / AL | 1.21 | 0.19 | 30.06 | 1.21 | 0.316 |
| AM | 6.0 | PL / PM | 1.11 | 0.18 | 27.18 | 0.65 | 0.659 |
| AM | 6.0 | AL / PM | 0.92 | 0.11 | 157.1 | -0.72 | 0.604 |
| AM | 8.0 | SP / AM | 0.45 | 0.07 | 31.36 | -5.02 | **< 0.001** |
| AM | 8.0 | SP / PL | 0.56 | 0.10 | 24.36 | -3.36 | **0.005** |
| AM | 8.0 | SP / AL | 1.00 | 0.17 | 26.17 | -0.02 | 1.000 |
| AM | 8.0 | SP / PM | 1.04 | 0.21 | 19.08 | 0.22 | 0.943 |
| AM | 8.0 | AM / PL | 1.24 | 0.16 | 72.48 | 1.63 | 0.155 |
| AM | 8.0 | AM / AL | 2.20 | 0.39 | 23.55 | 4.47 | **< 0.001** |
| AM | 8.0 | AM / PM | 2.30 | 0.43 | 21.37 | 4.50 | **< 0.001** |
| AM | 8.0 | PL / AL | 1.77 | 0.28 | 29.86 | 3.61 | **0.002** |
| AM | 8.0 | PL / PM | 1.86 | 0.31 | 27.07 | 3.77 | **0.002** |
| AM | 8.0 | AL / PM | 1.05 | 0.12 | 157.71 | 0.40 | 0.828 |
| PL | 0.5 | SP / AM | 3.02 | 0.47 | 30.99 | 7.03 | **< 0.001** |
| PL | 0.5 | SP / PL | 6.39 | 1.10 | 24.74 | 10.75 | **< 0.001** |
| PL | 0.5 | SP / AL | 2.8 | 0.47 | 26.49 | 6.17 | **< 0.001** |
| PL | 0.5 | SP / PM | 3.46 | 0.68 | 19.14 | 6.32 | **< 0.001** |
| PL | 0.5 | AM / PL | 2.12 | 0.28 | 71.75 | 5.74 | **< 0.001** |
| PL | 0.5 | AM / AL | 0.93 | 0.16 | 23.33 | -0.42 | 0.822 |
| PL | 0.5 | AM / PM | 1.15 | 0.21 | 21.05 | 0.75 | 0.597 |
| PL | 0.5 | PL / AL | 0.44 | 0.07 | 30.4 | -5.17 | **< 0.001** |
| PL | 0.5 | PL / PM | 0.54 | 0.09 | 27.3 | -3.71 | **0.002** |
| PL | 0.5 | AL / PM | 1.24 | 0.15 | 158.92 | 1.77 | 0.114 |
| PL | 2.5 | SP / AM | 2.40 | 0.38 | 31.28 | 5.55 | **< 0.001** |
| PL | 2.5 | SP / PL | 3.28 | 0.56 | 24.5 | 6.91 | **< 0.001** |
| PL | 2.5 | SP / AL | 3.18 | 0.53 | 26.44 | 6.94 | **< 0.001** |
| PL | 2.5 | SP / PM | 3.12 | 0.61 | 19.14 | 5.79 | **< 0.001** |
| PL | 2.5 | AM / PL | 1.37 | 0.18 | 71.51 | 2.41 | **0.030** |
| PL | 2.5 | AM / AL | 1.33 | 0.23 | 23.46 | 1.61 | 0.170 |
| PL | 2.5 | AM / PM | 1.30 | 0.24 | 21.19 | 1.42 | 0.233 |
| PL | 2.5 | PL / AL | 0.97 | 0.15 | 29.99 | -0.2 | 0.958 |
| PL | 2.5 | PL / PM | 0.95 | 0.16 | 27.01 | -0.31 | 0.885 |
| PL | 2.5 | AL / PM | 0.98 | 0.12 | 158.32 | -0.17 | 0.969 |
| PL | 6.0 | SP / AM | 0.58 | 0.09 | 31.35 | -3.44 | **0.003** |
| PL | 6.0 | SP / PL | 0.95 | 0.16 | 24.45 | -0.31 | 0.885 |
| PL | 6.0 | SP / AL | 1.03 | 0.17 | 26.38 | 0.17 | 0.969 |
| PL | 6.0 | SP / PM | 0.95 | 0.19 | 19.09 | -0.25 | 0.931 |
| PL | 6.0 | AM / PL | 1.63 | 0.21 | 71.51 | 3.73 | **0.001** |
| PL | 6.0 | AM / AL | 1.77 | 0.31 | 23.46 | 3.24 | **0.006** |
| PL | 6.0 | AM / PM | 1.64 | 0.30 | 21.15 | 2.66 | **0.024** |
| PL | 6.0 | PL / AL | 1.09 | 0.17 | 29.86 | 0.52 | 0.750 |
| PL | 6.0 | PL / PM | 1.01 | 0.17 | 26.84 | 0.03 | 1.000 |
| PL | 6.0 | AL / PM | 0.93 | 0.11 | 156.49 | -0.65 | 0.659 |
| PL | 8.0 | SP / AM | 0.43 | 0.07 | 31.42 | -5.32 | **< 0.001** |
| PL | 8.0 | SP / PL | 0.71 | 0.12 | 24.69 | -2.01 | 0.084 |
| PL | 8.0 | SP / AL | 0.99 | 0.17 | 26.49 | -0.04 | 1.000 |
| PL | 8.0 | SP / PM | 1.09 | 0.21 | 19.14 | 0.46 | 0.797 |
| PL | 8.0 | AM / PL | 1.64 | 0.22 | 72.94 | 3.76 | **0.001** |
| PL | 8.0 | AM / AL | 2.30 | 0.40 | 23.59 | 4.72 | **< 0.001** |
| PL | 8.0 | AM / PM | 2.53 | 0.47 | 21.26 | 5.02 | **< 0.001** |
| PL | 8.0 | PL / AL | 1.40 | 0.22 | 30.33 | 2.13 | 0.064 |
| PL | 8.0 | PL / PM | 1.55 | 0.26 | 27.24 | 2.64 | **0.022** |
| PL | 8.0 | AL / PM | 1.10 | 0.13 | 158.93 | 0.81 | 0.544 |

**Supplementary Table 3:** Multiple comparisons between test-stimulus orientations (TSO). The results are presented as follows, for a fixed conditioning-stimulus orientation (CSO) and interstimulus interval (ISI), we compute the motor evoked potential (MEP) amplitude ratio (MEP ratio) between the tested TSO. Each comparison has a standard error (SE), degrees of freedom (DoF), *t*-ratio, and *p*-value. Tested stimulus orientations were anteromedial (AM), posterolateral (PL), anterolateral (AL), and posteromedial (PM). *p*-values in bold are smaller than the statistical significance level of 0.05.

| CSO | ISI (ms) | MEP ratio<br>TSO (AM/PL) | SE | DoF | *t*-ratio | *p*-value |
| --- | --- | --- | --- | --- | --- | --- |
| SP | 0.5 | 1.88 | 0.29 | 39.59 | 4.15 | **< 0.001** |
| SP | 2.5 | 1.88 | 0.29 | 39.59 | 4.15 | **< 0.001** |
| SP | 6.0 | 1.88 | 0.29 | 39.59 | 4.15 | **< 0.001** |
| SP | 8.0 | 1.88 | 0.29 | 39.59 | 4.15 | **< 0.001** |
| AM | 0.5 | 0.59 | 0.09 | 39.39 | -3.51 | **0.002** |
| AM | 2.5 | 1.30 | 0.20 | 39.88 | 1.72 | 0.137 |
| AM | 6.0 | 1.47 | 0.23 | 40.27 | 2.52 | **0.026** |
| AM | 8.0 | 1.79 | 0.28 | 40.67 | 3.80 | **0.001** |
| PL | 0.5 | 1.82 | 0.28 | 39.88 | 3.92 | **0.001** |
| PL | 2.5 | 1.35 | 0.21 | 39.98 | 1.97 | 0.084 |
| PL | 6.0 | 1.95 | 0.30 | 39.68 | 4.39 | **< 0.001** |
| PL | 8.0 | 2.37 | 0.36 | 39.88 | 5.65 | **< 0.001** |
| AL | 0.5 | 2.56 | 0.39 | 39.78 | 6.15 | **< 0.001** |
| AL | 2.5 | 2.16 | 0.33 | 39.87 | 5.05 | **< 0.001** |
| AL | 6.0 | 1.75 | 0.27 | 39.39 | 3.67 | **0.001** |
| AL | 8.0 | 1.88 | 0.29 | 39.59 | 4.13 | **< 0.001** |
| PM | 0.5 | 2.50 | 0.38 | 39.49 | 6.01 | **< 0.001** |
| PM | 2.5 | 2.21 | 0.34 | 39.88 | 5.20 | **< 0.001** |
| PM | 6.0 | 1.77 | 0.27 | 39.30 | 3.74 | **0.001** |
| PM | 8.0 | 1.97 | 0.30 | 39.58 | 4.45 | **< 0.001** |

**Supplementary Table 4:** Multiple comparisons between interstimulus intervals (ISI). The results are presented as follows, for a fixed conditioning stimulus orientation (CSO) and test stimulus orientation (TSO), we compute the motor evoked potential latency difference between the tested ISI. Each comparison has a standard error (SE), degrees of freedom (DoF), *t*-ratio, and *p*-value. Tested stimulus orientations were anteromedial (AM), posterolateral (PL), anterolateral (AL), and posteromedial (PM); ISI is given in milliseconds. *p*-values in bold are smaller than the statistical significance level of 0.05.

| CSO | TSO | Comparison<br>ISI<sub>1</sub> (ms) – ISI<sub>2</sub> (ms) | Latency<br>difference (ms) | SE | DoF | *t*-ratio | *p*-value |
| --- | --- | --- | --- | --- | --- | --- | --- |
| AM | AM | 0.5 – 2.5 | 1.42 | 0.16 | 5659.67 | 9.12 | < **0.001** |
| AM | AM | 0.5 – 6.0 | 1.92 | 0.15 | 5505.14 | 12.99 | < **0.001** |
| AM | AM | 0.5 – 8.0 | 1.82 | 0.15 | 5454.97 | 12.32 | < **0.001** |
| AM | AM | 2.5 – 6.0 | 0.5 | 0.11 | 5640.95 | 4.54 | < **0.001** |
| AM | AM | 2.5 – 8.0 | 0.4 | 0.11 | 5597.93 | 3.65 | **0.001** |
| AM | AM | 6.0 – 8.0 | –0.1 | 0.09 | 5674.76 | –1.06 | 0.45 |
| PL | AM | 0.5 – 2.5 | 1.02 | 0.15 | 4992.93 | 6.81 | < **0.001** |
| PL | AM | 0.5 – 6.0 | 1.05 | 0.13 | 5186.69 | 8 | < **0.001** |
| PL | AM | 0.5 – 8.0 | 1.01 | 0.13 | 5225.58 | 7.78 | < **0.001** |
| PL | AM | 2.5 – 6.0 | 0.03 | 0.12 | 5007.23 | 0.23 | 0.943 |
| PL | AM | 2.5 – 8.0 | –0.01 | 0.12 | 4634.97 | –0.12 | 1.0 |
| PL | AM | 6.0 – 8.0 | –0.04 | 0.09 | 5681.71 | –0.45 | 0.811 |
| AL | AM | 0.5 – 2.5 | 0.01 | 0.11 | 5664.8 | 0.11 | 1.0 |
| AL | AM | 0.5 – 6.0 | 0.02 | 0.1 | 5645.93 | 0.2 | 0.962 |
| AL | AM | 0.5 – 8.0 | –0.03 | 0.1 | 5669.95 | –0.34 | 0.884 |
| AL | AM | 2.5 – 6.0 | 0.01 | 0.11 | 5467.64 | 0.08 | 1.0 |
| AL | AM | 2.5 – 8.0 | –0.05 | 0.11 | 5539.65 | –0.44 | 0.811 |
| AL | AM | 6.0 – 8.0 | –0.06 | 0.1 | 5674.72 | –0.57 | 0.747 |
| PM | AM | 0.5 – 2.5 | 0.09 | 0.12 | 5677.21 | 0.79 | 0.605 |
| PM | AM | 0.5 – 6.0 | 0.15 | 0.11 | 5394.38 | 1.39 | 0.288 |
| PM | AM | 0.5 – 8.0 | 0.14 | 0.11 | 5206.83 | 1.29 | 0.336 |
| PM | AM | 2.5 – 6.0 | 0.06 | 0.11 | 5428.48 | 0.56 | 0.747 |
| PM | AM | 2.5 – 8.0 | 0.05 | 0.11 | 5300.91 | 0.46 | 0.803 |
| PM | AM | 6.0 – 8.0 | –0.01 | 0.1 | 5678.89 | –0.1 | 1.0 |
| AM | PL | 0.5 – 2.5 | 0.71 | 0.14 | 5681.24 | 5.14 | < **0.001** |
| AM | PL | 0.5 – 6.0 | 1.02 | 0.13 | 5604.52 | 8.11 | < **0.001** |
| AM | PL | 0.5 – 8.0 | 0.78 | 0.13 | 5541.44 | 6.23 | < **0.001** |
| AM | PL | 2.5 – 6.0 | 0.31 | 0.12 | 5599.52 | 2.61 | **0.021** |
| AM | PL | 2.5 – 8.0 | 0.07 | 0.12 | 5533.62 | 0.56 | 0.747 |
| AM | PL | 6.0 – 8.0 | –0.24 | 0.1 | 5677.83 | –2.47 | **0.030** |
| PL | PL | 0.5 – 2.5 | 1.14 | 0.18 | 5672.35 | 6.16 | < **0.001** |
| PL | PL | 0.5 – 6.0 | 1.36 | 0.17 | 5529.51 | 7.96 | < **0.001** |
| PL | PL | 0.5 – 8.0 | 0.99 | 0.17 | 5452.67 | 5.8 | < **0.001** |
| PL | PL | 2.5 – 6.0 | 0.22 | 0.13 | 5491.87 | 1.7 | 0.162 |
| PL | PL | 2.5 – 8.0 | –0.15 | 0.13 | 5347.54 | –1.13 | 0.416 |
| PL | PL | 6.0 – 8.0 | –0.37 | 0.1 | 5681.94 | –3.6 | **0.001** |
| AL | PL | 0.5 – 2.5 | 0.18 | 0.14 | 5680.02 | 1.28 | 0.336 |
| AL | PL | 0.5 – 6.0 | 0.35 | 0.13 | 5602.57 | 2.75 | **0.014** |
| AL | PL | 0.5 – 8.0 | 0.14 | 0.13 | 5502.02 | 1.12 | 0.417 |
| AL | PL | 2.5 – 6.0 | 0.16 | 0.13 | 5462.02 | 1.23 | 0.362 |
| AL | PL | 2.5 – 8.0 | -0.04 | 0.13 | 5363.73 | -0.33 | 0.887 |
| AL | PL | 6.0 – 8.0 | -0.21 | 0.11 | 5680.36 | -1.9 | 0.111 |
| PM | PL | 0.5 – 2.5 | 0.13 | 0.15 | 5622.91 | 0.9 | 0.532 |
| PM | PL | 0.5 – 6.0 | 0.23 | 0.13 | 4943.53 | 1.79 | 0.138 |
| PM | PL | 0.5 – 8.0 | 0.17 | 0.13 | 4832.83 | 1.31 | 0.329 |
| PM | PL | 2.5 – 6.0 | 0.1 | 0.13 | 5189.32 | 0.78 | 0.605 |
| PM | PL | 2.5 – 8.0 | 0.04 | 0.13 | 5118.95 | 0.31 | 0.893 |
| PM | PL | 6.0 – 8.0 | -0.06 | 0.11 | 5676.86 | -0.57 | 0.747 |

**Supplementary Table 5:** Multiple comparisons between conditioning stimulus orientations (CSO). The results are presented as follows, for a fixed test stimulus orientation (TSO) and interstimulus interval (ISI), we compute the motor evoked potential latency difference between the tested CSO. Each comparison has a standard error (SE), degrees of freedom (DoF), *t*-ratio, and *p*-value. Tested stimulus orientations were anteromedial (AM), posterolateral (PL), anterolateral (AL), and posteromedial (PM); ISI is given in milliseconds. *p*-values in bold are smaller than the statistical significance level of 0.05.

| TSO | ISI (ms) | Comparison CSO<sub>1</sub> – CSO<sub>2</sub> | Latency difference (ms) | SE | DoF | *t*-ratio | *p*-value |
| --- | --- | --- | --- | --- | --- | --- | --- |
| AM | 0.5 | SP – AM | -0.80 | 0.21 | 32.45 | -3.75 | **0.002** |
| AM | 0.5 | SP – PL | -0.26 | 0.17 | 40.78 | -1.47 | 0.267 |
| AM | 0.5 | SP – AL | 0.10 | 0.13 | 53.28 | 0.79 | 0.605 |
| AM | 0.5 | SP – PM | 0.030 | 0.15 | 37.17 | 0.18 | 0.975 |
| AM | 0.5 | AM – PL | 0.54 | 0.19 | 239.57 | 2.92 | **0.009** |
| AM | 0.5 | AM – AL | 0.90 | 0.18 | 75.00 | 4.88 | **< 0.001** |
| AM | 0.5 | AM – PM | 0.82 | 0.17 | 149.69 | 4.71 | **< 0.001** |
| AM | 0.5 | PL – AL | 0.35 | 0.16 | 76.23 | 2.21 | 0.064 |
| AM | 0.5 | PL – PM | 0.28 | 0.15 | 165.62 | 1.84 | 0.130 |
| AM | 0.5 | AL – PM | -0.07 | 0.12 | 192.91 | -0.60 | 0.736 |
| AM | 2.5 | SP – AM | 0.62 | 0.19 | 20.21 | 3.28 | **0.009** |
| AM | 2.5 | SP – PL | 0.76 | 0.17 | 34.84 | 4.56 | **< 0.001** |
| AM | 2.5 | SP – AL | 0.11 | 0.13 | 59.00 | 0.86 | 0.566 |
| AM | 2.5 | SP – PM | 0.12 | 0.14 | 35.52 | 0.82 | 0.597 |
| AM | 2.5 | AM – PL | 0.14 | 0.15 | 106.02 | 0.96 | 0.502 |
| AM | 2.5 | AM – AL | -0.51 | 0.16 | 41.43 | -3.21 | **0.007** |
| AM | 2.5 | AM – PM | -0.50 | 0.14 | 69.36 | -3.51 | **0.002** |
| AM | 2.5 | PL – AL | -0.65 | 0.16 | 67.88 | -4.16 | **< 0.001** |
| AM | 2.5 | PL – PM | -0.65 | 0.14 | 127.27 | -4.47 | **< 0.001** |
| AM | 2.5 | AL – PM | 0.01 | 0.12 | 193.37 | 0.05 | 1.0 |
| AM | 6.0 | SP – AM | 1.12 | 0.18 | 16.79 | 6.20 | **< 0.001** |
| AM | 6.0 | SP – PL | 0.79 | 0.15 | 22.88 | 5.30 | **< 0.001** |
| AM | 6.0 | SP – AL | 0.12 | 0.12 | 45.89 | 0.99 | 0.494 |
| AM | 6.0 | SP – PM | 0.18 | 0.14 | 28.76 | 1.31 | 0.336 |
| AM | 6.0 | AM – PL | -0.32 | 0.12 | 44.50 | -2.76 | **0.020** |
| AM | 6.0 | AM – AL | -1.00 | 0.14 | 26.25 | -7.11 | **< 0.001** |
| AM | 6.0 | AM – PM | -0.94 | 0.12 | 39.04 | -7.69 | **< 0.001** |
| AM | 6.0 | PL – AL | -0.67 | 0.13 | 33.70 | -5.18 | **< 0.001** |
| AM | 6.0 | PL – PM | -0.61 | 0.11 | 56.21 | -5.39 | **< 0.001** |
| AM | 6.0 | AL – PM | 0.06 | 0.10 | 119.27 | 0.56 | 0.747 |
| AM | 8.0 | SP – AM | 1.02 | 0.18 | 16.68 | 5.66 | **< 0.001** |
| AM | 8.0 | SP – PL | 0.75 | 0.15 | 22.00 | 5.07 | **< 0.001** |
| AM | 8.0 | SP – AL | 0.06 | 0.12 | 45.44 | 0.53 | 0.765 |
| AM | 8.0 | SP – PM | 0.17 | 0.14 | 29.49 | 1.23 | 0.373 |
| AM | 8.0 | AM – PL | -0.27 | 0.12 | 41.22 | -2.33 | 0.053 |
| AM | 8.0 | AM – AL | -0.95 | 0.14 | 25.75 | -6.84 | **< 0.001** |
| AM | 8.0 | AM – PM | -0.85 | 0.12 | 39.73 | -6.93 | **< 0.001** |
| AM | 8.0 | PL – AL | -0.69 | 0.13 | 31.70 | -5.37 | **< 0.001** |
| AM | 8.0 | PL – PM | -0.58 | 0.11 | 54.70 | -5.14 | **< 0.001** |
| AM | 8.0 | AL – PM | 0.10 | 0.11 | 122.06 | 0.99 | 0.493 |
| PL | 0.5 | SP – AM | 0.22 | 0.20 | 26.18 | 1.09 | 0.449 |
| PL | 0.5 | SP – PL | -0.37 | 0.21 | 82.50 | -1.76 | 0.152 |
| PL | 0.5 | SP – AL | -0.30 | 0.15 | 94.68 | -2.01 | 0.097 |
| PL | 0.5 | SP – PM | -0.22 | 0.17 | 58.82 | -1.32 | 0.332 |
| PL | 0.5 | AM – PL | -0.59 | 0.20 | 313.63 | -2.94 | **0.009** |
| PL | 0.5 | AM – AL | -0.51 | 0.18 | 65.27 | -2.90 | 0.012 |
| PL | 0.5 | AM – PM | -0.44 | 0.17 | 129.9 | -2.59 | **0.024** |
| PL | 0.5 | PL – AL | 0.07 | 0.20 | 188.16 | 0.35 | 0.881 |
| PL | 0.5 | PL – PM | 0.15 | 0.20 | 406.34 | 0.75 | 0.625 |
| PL | 0.5 | AL – PM | 0.08 | 0.15 | 372.86 | 0.51 | 0.770 |
| PL | 2.5 | SP – AM | 0.93 | 0.20 | 23.57 | 4.75 | **< 0.001** |
| PL | 2.5 | SP – PL | 0.77 | 0.18 | 43.28 | 4.36 | **< 0.001** |
| PL | 2.5 | SP – AL | -0.11 | 0.15 | 106.98 | -0.73 | 0.64 |
| PL | 2.5 | SP – PM | -0.08 | 0.16 | 57.15 | -0.52 | 0.77 |
| PL | 2.5 | AM – PL | -0.16 | 0.16 | 132.85 | -1.02 | 0.476 |
| PL | 2.5 | AM – AL | -1.05 | 0.18 | 62.48 | -5.93 | **< 0.001** |
| PL | 2.5 | AM – PM | -1.02 | 0.16 | 108.46 | -6.31 | **< 0.001** |
| PL | 2.5 | PL – AL | -0.88 | 0.18 | 104.52 | -5.02 | **< 0.001** |
| PL | 2.5 | PL – PM | -0.86 | 0.16 | 195.45 | -5.21 | **< 0.001** |
| PL | 2.5 | AL – PM | 0.03 | 0.16 | 376.68 | 0.17 | 0.975 |
| PL | 6.0 | SP – AM | 1.24 | 0.19 | 18.99 | 6.67 | **< 0.001** |
| PL | 6.0 | SP – PL | 0.99 | 0.16 | 27.98 | 6.30 | **< 0.001** |
| PL | 6.0 | SP – AL | 0.05 | 0.13 | 63.29 | 0.39 | 0.856 |
| PL | 6.0 | SP – PM | 0.02 | 0.15 | 36.62 | 0.11 | 1.0 |
| PL | 6.0 | AM – PL | -0.25 | 0.12 | 53.02 | -2.00 | 0.098 |
| PL | 6.0 | AM – AL | -1.19 | 0.15 | 30.75 | -8.14 | **< 0.001** |
| PL | 6.0 | AM – PM | -1.22 | 0.13 | 46.12 | -9.60 | **< 0.001** |
| PL | 6.0 | PL – AL | -0.94 | 0.14 | 41.96 | -6.86 | **< 0.001** |
| PL | 6.0 | PL – PM | -0.98 | 0.12 | 70.95 | -8.06 | **< 0.001** |
| PL | 6.0 | AL – PM | -0.03 | 0.11 | 160.17 | -0.30 | 0.893 |
| PL | 8.0 | SP – AM | 1.0 | 0.19 | 18.73 | 5.40 | **< 0.001** |
| PL | 8.0 | SP – PL | 0.62 | 0.16 | 27.11 | 3.99 | **0.001** |
| PL | 8.0 | SP – AL | -0.15 | 0.13 | 62.75 | -1.17 | 0.398 |
| PL | 8.0 | SP – PM | -0.04 | 0.15 | 37.88 | -0.30 | 0.893 |
| PL | 8.0 | AM – PL | -0.38 | 0.12 | 49.02 | -3.11 | **0.008** |
| PL | 8.0 | AM – AL | -1.15 | 0.14 | 29.84 | -7.96 | **< 0.001** |
| PL | 8.0 | AM – PM | -1.04 | 0.13 | 46.85 | -8.15 | **< 0.001** |
| PL | 8.0 | PL – AL | -0.78 | 0.14 | 40.01 | -5.74 | **< 0.001** |
| PL | 8.0 | PL – PM | -0.67 | 0.12 | 70.95 | -5.51 | **< 0.001** |
| PL | 8.0 | AL – PM | 0.11 | 0.12 | 166.16 | 0.95 | 0.502 |

**Supplementary Table 6:** Multiple comparisons between test stimulus orientations (TSO). The results are presented as follows, for a fixed conditioning stimulus orientation (CSO) and interstimulus interval (ISI), we compute the motor evoked potential latency difference between the tested TSO. Each comparison has a standard error (SE), degrees of freedom (DoF), *t*-ratio, and *p*-value. Tested stimulus orientations were anteromedial (AM), posterolateral (PL), anterolateral (AL), and posteromedial (PM); ISI is given in milliseconds. *p*-values in bold are smaller than the statistical significance level of 0.05.

| CSO | ISI (ms) | Latency difference TSO (AM – PL) | SE | DoF | *t*-ratio | *p*-value |
| --- | --- | --- | --- | --- | --- | --- |
| SP | 0.5 | -0.23 | 0.11 | 174.04 | -1.98 | 0.097 |
| SP | 2.5 | -0.23 | 0.11 | 174.04 | -1.98 | 0.097 |
| SP | 6.0 | -0.23 | 0.11 | 174.04 | -1.98 | 0.097 |
| SP | 8.0 | -0.23 | 0.11 | 174.04 | -1.98 | 0.097 |
| AM | 0.5 | 0.79 | 0.17 | 802.13 | 4.55 | **< 0.001** |
| AM | 2.5 | 0.09 | 0.14 | 333.4 | 0.65 | 0.704 |
| AM | 6.0 | -0.1 | 0.11 | 141.76 | -0.96 | 0.502 |
| AM | 8.0 | -0.25 | 0.11 | 133.94 | -2.31 | **0.049** |
| PL | 0.5 | -0.34 | 0.19 | 1164.94 | -1.73 | 0.153 |
| PL | 2.5 | -0.22 | 0.15 | 504.06 | -1.43 | 0.270 |
| PL | 6.0 | -0.03 | 0.11 | 161.84 | -0.23 | 0.943 |
| PL | 8.0 | -0.35 | 0.11 | 143.18 | -3.26 | **0.004** |
| AL | 0.5 | -0.62 | 0.14 | 329.16 | -4.59 | **< 0.001** |
| AL | 2.5 | -0.45 | 0.14 | 402.88 | -3.13 | **0.005** |
| AL | 6.0 | -0.3 | 0.11 | 176.98 | -2.58 | **0.024** |
| AL | 8.0 | -0.45 | 0.11 | 173.36 | -3.91 | **< 0.001** |
| PM | 0.5 | -0.47 | 0.14 | 408.72 | -3.26 | **0.003** |
| PM | 2.5 | -0.43 | 0.14 | 377.36 | -3.04 | **0.007** |
| PM | 6.0 | -0.39 | 0.11 | 162.25 | -3.47 | **0.002** |
| PM | 8.0 | -0.44 | 0.11 | 177.01 | -3.84 | **0.001** |

